# Dissociable contributions of the amygdala to the immediate and delayed effects of emotional arousal on memory

**DOI:** 10.1101/269050

**Authors:** Dirk Schümann, Tobias Sommer

## Abstract

Emotionally arousal enhances memory encoding and consolidation leading to better immediate and delayed memory. Although the central noradrenergic system and the amygdala play critical roles in both effects of emotional arousal, we have recently shown that these effects are at least partly independent of each other, suggesting distinct underlying neural mechanisms. Here we aim to dissociate the neural substrates of both effects in 70 female participants using an emotional memory paradigm to investigate how neural activity, as measured by fMRI, and a polymorphism in the α_2B_-noradrenoceptor vary for these effects. To also test whether the immediate and delayed effects of emotional arousal on memory are stable traits, we invited back participants who were a part of a large-scale behavioral memory study about 3.5 years ago. We replicated the low correlation of the immediate and delayed emotional enhancement of memory across participants (r = 0.16) and observed, moreover, that only the delayed effect was, to some degree, stable over time (r = 0.23). Bilateral amygdala activity, as well as its coupling with the visual cortex and the fusiform gyrus, was related to the preferential encoding of emotional stimuli, which is consistent with affect-biased attention. Moreover, the adrenoceptor genotype modulated the bilateral amygdala activity associated with this effect. The left amygdala and its coupling with the hippocampus was specifically associated with the more efficient consolidation of emotional stimuli, which is consistent with amygdalar modulation of hippocampal consolidation.

## Introduction

Improved memory for emotionally arousing compared to neutral information is driven by enhanced processing and encoding as well as by improved subsequent consolidation (Ritchey et al. 2008; Hamann 2001). Preferential encoding of emotional stimuli results immediately in enhanced memory and is based on their tendency to attract greater attention, resulting in deeper processing (Kensinger and Corkin 2004; Kang et al. 2014). This affect-biased attention, attributed to amygdala and central noradrenergic system activity, has been suggested to increase signal-to-noise ratio in sensory areas and synaptic plasticity in the hippocampus (Mather et al. 2015; Markovic et al. 2014; Hagena et al. 2016). The delayed effect of emotional arousal on consolidation is also mediated by the amygdala and the central noradrenergic system (McIntyre et al. 2012; Manns and Bass 2016; Inman et al. 2017). This delayed effect of arousal on consolidation can be explained by the tagging of active synapses during the processing of emotional stimuli that results in the subsequent transformation of early in persistent late long-term potentiation and hence improve consolidation (Bergado et al. 2011; McIntyre et al. 2012).

Given the critical roles played by the central noradrenergic system and the amygdala on the effects of emotional arousal on encoding and consolidation, it is plausible that the same core mechanisms that result downstream in the immediate emotional enhancement of memory (iEEM) also give rise to the delayed emotional enhancement of memories (dEEM). However, in a recent large-scale behavioral-genetic study (n = 690), we observed that the magnitude of both effects correlated only weakly with each other across participants (r = 0.14) (Schümann et al. under review). For instance, in some participants, emotional arousal only enhanced encoding (better memory for emotional items 10 minutes after encoding) but did not result in more efficient consolidation (memory for emotional and neutral items were similar 20 hours after encoding), whereas others showed the opposite pattern, i.e. only an effect on consolidation (better memory for emotionally arousing than neutral items after 20 hours but not immediately). The relative independence of these two effects from each other suggests at least partly distinct underlying processes and neurobiological substrates.

Three previous fMRI studies had aimed to disentangle the neural substrates of iEEM and dEEM (Ritchey et al. 2008; Mackiewicz et al. 2006; Mickley Steinmetz et al. 2012), but results are not fully consistent and open questions remain. The first study correlated mean activity of the amygdala during processing of negative compared to neutral pictures (i.e. the main effect of emotional arousal) across participants with memory performance in an immediate and a delayed recognition test (Mackiewicz et al. 2006). Bilateral dorsal amygdala was related to iEEM, the left ventral amygdala in women and the right in men to dEEM. Only the other two studies used subsequent memory effect (SME) analyses (Dolcos et al. 2012) to identify areas where trial-wise neural activity during encoding was correlated with iEEM or dEEM. The greater persistence of emotional memories, i.e. that dEEM is often behaviorally greater than that of iEEM, was not related to greater activity in the amygdala but increased coupling of the amygdala and the parahippocampal cortex (Ritchey et al. 2008). This study did not aim to identify which brain areas were more involved in iEEM. The third study tested the hypothesis that the subsequent memory effect for emotional items is more delay-invariant than for neutral items (Mickley Steinmetz et al. 2012). More cortical areas correlated with immediate than delayed subsequent memory, only for neutral – not negative – items, as predicted. However, no activity related to either iEEM or dEEM in the amygdala was observed.

In the study we present here, we aimed to complement the findings of these previous studies by directly comparing the neural correlates of iEEM and dEEM, i.e. the interaction of immediate and delayed SMEs, respectively. The difference can be considered in terms of affect-biased attention (i.e. iEEM) versus emotional synaptic tagging (i.e. dEEM). In addition, we aimed to contrast the functional coupling of the amygdala with other brain areas related to iEEM and dEEM, given that affect-biased attention and emotional synaptic tagging rely on amygdala modulation of different brain areas. Such direct comparisons of iEEM and dEEM is challenging because many emotionally arousing items involve both enhanced attention and synaptic tagging, though our previous study suggests that both processes are relatively independent of each other (Schümann et al. under review). However, these items cannot be identified and then excluded from the analyses because each item can be tested only once, immediately or delayed (otherwise the second test would be confounded by the first). These confounded items can only be treated as noise when one aims to separate the neural correlates of the iEEM and dEEM during the initial processing. To overcome this limitation inherent to the nature of EEM, we recruited a relatively large sample of 70 participants in an attempt to increase statistical power needed to detect activity uniquely associated with iEEM or with dEEM. Only female participants were recruited since lateralized amygdala activity related to the dEEM depending on sex has been observed (Cahill et al. 2004; Mackiewicz et al. 2006).

In our previous large-scale behavioral study, we had tested whether iEEM and dEEM’s independence of each other could be explained by the differential involvement of the adrenoceptor subtypes, which vary in affinity, action and expression pattern (Hein 2006; Luhrs et al. 2016). Participants were genotyped for a polymorphism in the gene that codes for the α_2B_-noradrenergic receptor, a polymorphism that has been associated with greater perceived vividness of emotional stimuli, affect-biased attention, and iEEM in free recall (de Quervain et al. 2007; Rasch et al. 2009; Todd et al. 2015, 2013). However, we did not find any relationship between the polymorphism and iEEM or dEEM in behavior (Schümann et al. under review). Given that differences in neural activity are not necessarily behaviorally observable, we invited back a subset of the genotyped participants from our previous large-scale behavioral study in order to investigate whether α_2B_-noradrenoceptor polymorphism is associated with differences in neural activity between iEEM and dEEM. This also enabled us to investigate to what degree iEEM and dEEM are stable personality traits, which would in turn point to genetic influences.

## Methods

### Participants

70 female participants (mean age 28 years, range 22-38 years) who had participated in the large behavioral study 3 years 5 months earlier on average (range 1y4m to 5y10m) were recruited. Data acquisition failed for one participant due to equipment malfunction, leaving a sample of 69. The study was approved by the local ethics committee of the Hamburg Board of Physicians in Germany. All participants signed informed consent and were paid €10 per hour for the participation.

### Emotional Memory Paradigm

Participants studied 80 neutral and 80 negatively arousing scenes in the MRI scanner. Scenes were drawn from a set of 160 neutral and 160 negative scenes. Only negative, that is no positive, pictures were contrasted with neutral ones to increase statistical power as negative and positive EEMs partially rely on distinct mechanisms and neural substrates (Mickley Steinmetz et al. 2010; Mickley Steinmetz and Kensinger 2009; Talmi et al. 2007). Each trial consisted of picture presentation (2 s) followed by rating the suitability of the picture for a magazine like National Geographic (2 s), an active baseline task (pointing arrows, 4 s) and a jittered fixation period (0.95 to 3.05 s), resulting in trial lengths of about 9 to 12 s. The trial structure, including the active baseline task, was very similar to what was used in our previous large-scale behavioral study. The presentation rate of the pictures was slow enough to allow a valid psychophysiological interaction (PPI) analyses (Gitelman et al. 2003; Friston et al. 1997). At the end of this encoding period in the MRI scanner, participants completed 3 minutes of a mental rotation task to clear working memory outside of the scanner. They then completed a recognition test with a 6-point confidence scale for half of the pictures, randomly intermixed with the same number of unseen pictures as lures. A second recognition test for the remaining pictures followed about 24 hours later. At the conclusion of the study, participants rated the subjective valence and arousal of the pictures on a 9-step Self-Assessment Manikin (Bradley and Lang 1994), which confirmed that the negative pictures were experienced as more negative and more arousing than the neutral ones, with mean valence for negative pictures 2.9 (SD 0.7), neutral 6.6 (SD 1.1), t(68) = −24,43, p < 0.0001; mean arousal for negative pictures 6.3 (SD 1.2), neutral 2.6 (SD 1.1), t(68) = 23,00, p < 0.0001.

### Neuroimaging

Event-related functional whole brain MRI was performed on a 3 Tesla system (Siemens Trio) with a T2*-weighted echo planar imaging sequence in 38 contiguous axial slices (3-mm thickness with 1-mm gap; TR 2.21 s; TE 30 ms; flip angle 80°; FOV 216 × 216; matrix 72 × 72). For spatial normalization, a high-resolution T1-weighted structural MR image was acquired with a 3D MPRAGE sequence (TR 2300 ms, TE 2.89 ms, flip angle 9°, 1-mm slices, FOV 256 × 192; 240 slices).

Statistical Parametric Mapping (SPM12; Wellcome Department of Imaging Neuroscience, London, UK) was used to preprocess and analyze the fMRI data. The fMRI data were slice-time corrected, realigned and corrected for susceptibility-by-movement artifacts using the realign and unwarp function as implemented in SPM12. T1 images were then co-registered to the individual mean functional image, segmented and normalized using the DARTEL toolbox to create individual flow fields. Flow fields were also used for normalizing the functional images to MNI-space. Finally, images were smoothed with a full-width half-maximum Gaussian kernel of 8 mm.

Each participant’s fMRI data was modeled using a general linear model (GLM) with 2 emotional arousal (neutral vs. negative) × 2 subsequent memory (subsequent hits vs. misses) × 2 delay (immediate vs. delayed) regressors that were created by convolving the respective onset vectors (i.e. in an event-related manner) with the canonical hemodynamic response function. To remove movement-related artifacts, for images with movements greater than 0.2 mm/TR nuisance regressors were created. The time series were corrected for baseline drifts by applying a high-pass filter (128 s) and for serial dependency using an AR(1) autocorrelation model. The contrast images corresponding to the regressors of interest for all participants were then included in a group-level GLM. To identify areas where activity correlated with iEEM, the interaction of encoding success (SME) and emotional arousal was computed for the immediately tested images. For dEEM, the same interaction was computed but for images tested after 24 hours. In order to identify areas predominantly involved in the immediate vs. delayed EEM the particular contrast was exclusively masked (using a liberal threshold of p < 0.05 uncorrected) by the other. This masking procedure rather than more rigid statistical interactions was used to deal with the complication that both processes are innate to the affective processing of many pictures, making their separation difficult even with large samples.

PPI analyses were conducted, as implemented in SPM12 (Friston et al. 1997), to assess differences in the functional coupling of the amygdala between successful memory formation of negative and neutral pictures. In particular, we tested with which brain areas the part of the amygdala involved in emotional processing (main effect of emotion, xyz = [−20 −6 −14], Z = inf, 243 voxels) was more strongly coupled for negative than neutral pictures during successful immediate vs. delayed memory formation. Again an exclusive masking procedure (p < 0.05, uncorrected) was applied in order to identify areas more strongly coupled during immediate than delayed EEM and vice versa.

To assess genotype-dependent differences in brain activity, we contrasted deletion carriers (homo-and heterozygote) vs. homozygote wildtype for the α_2B_-noradrenergic receptor polymorphism by adding genotype group as a factor to the second level analysis (Rasch et al. 2009; Todd et al. 2015; Urner et al. 2011). Interactions of genotype with the main effect of emotion as well as with the interaction contrasts representing iEEM and dEEM were computed.

All voxel coordinates are given in MNI-space..Results of all analyses were considered significant at p < 0.05, peak voxel family-wise error corrected for multiple comparisons within previously reported regions involved in EEM, where the amygdala, hippocampus and visual areas were most relevant for the hypotheses of the current study (Kensinger and Schacter 2007; Ritchey et al. 2011). Anatomical masks of the Automatic Anatomic Labeling toolbox were used for small volume correction (Tzourio-Mazoyer et al. 2002).

## Results

### Behavioral Results

Corrected hit rates (hit rate – false alarm rate) were higher for negative than for neutral stimuli at both retention intervals (Figure 1, left panel). A GLM with within-subject factors *retention interval* and *valence* revealed main effects of *retention interval*, F(1,68) = 306.98, p < 0.0001, η^2^ = 0.24, and *valence*, F(1,68) = 83.08, p < 0.0001, η^2^ = 0.07, and an interaction between these two factors, F(1,68) = 29.32, p < 0.0001, η^2^ = 0.01, indicating that the effect of emotional arousal was larger after a consolidation delay. There was no correlation between delayed (dEEM) and immediate (iEEM) effects of emotional arousal: r = 0.16 (p = 0.20; Figure 1, middle panel), which replicates our earlier finding. To give a full picture of the behavioral results, we computed in addition to the corrected hit rates also d-prime, response bias (c), and - as a pure measure of accuracy (Dougal and Rotello 2007) - the area under the curve (AUC) of the ROC-curves based on the confidence ratings (Table 1). Finally, to assess the contribution of recollection and familiarity to the EEMs the dual process model of recognition memory was fit to the individual data using the ROC toolbox (Koen et al. 2016).

**Figure 1:**
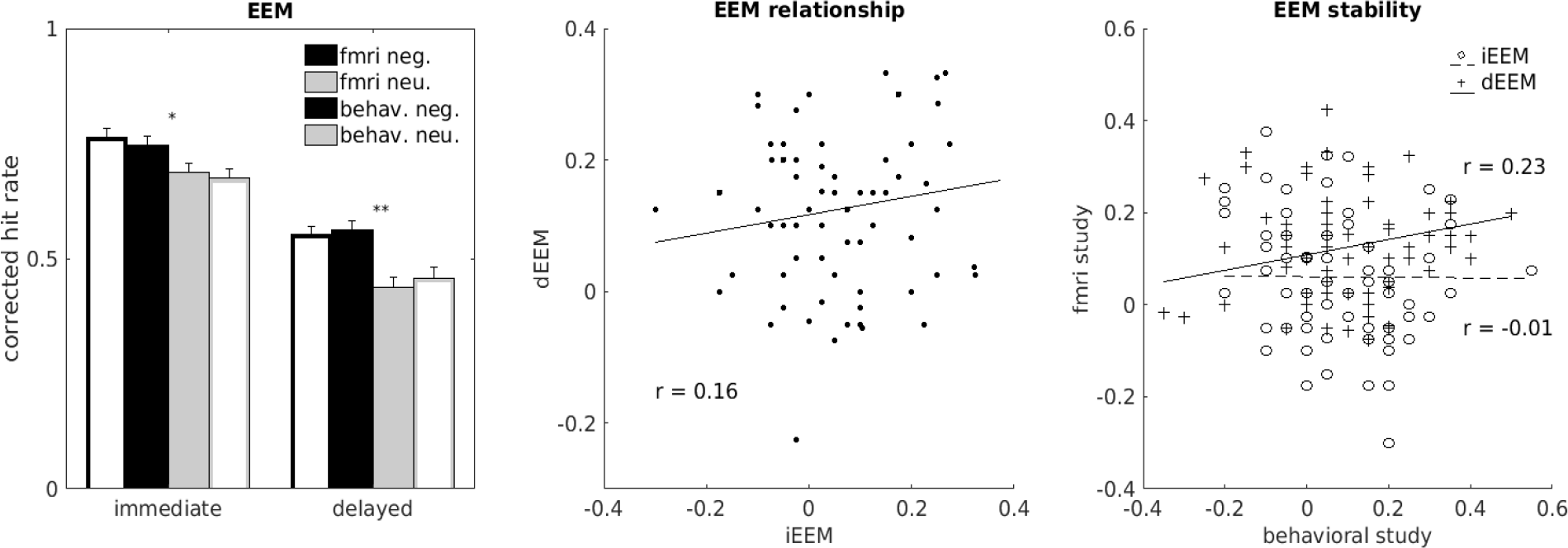
Behavioral results. *Left panel*: Corrected hit rates in the immediate and delayed memory test for negative (black) and neutral (gray) pictures from the current study and from the previous large-scale behavioral study of the same participants (solid bars – current fMRI study, outlined bars – large-scale behavioral study). *Middle panel*: Correlations between the immediate and delayed emotional enhancement of memory (iEEM and dEEM, respectively) across participants in the current study. Exclusion of the outlier changes correlation to r = 0.14. *Right panel*: Correlations of iEEMs, respectively dEEMs from the current study and from the previous large-scale behavioral study of the same participants.

**Figure 2:**
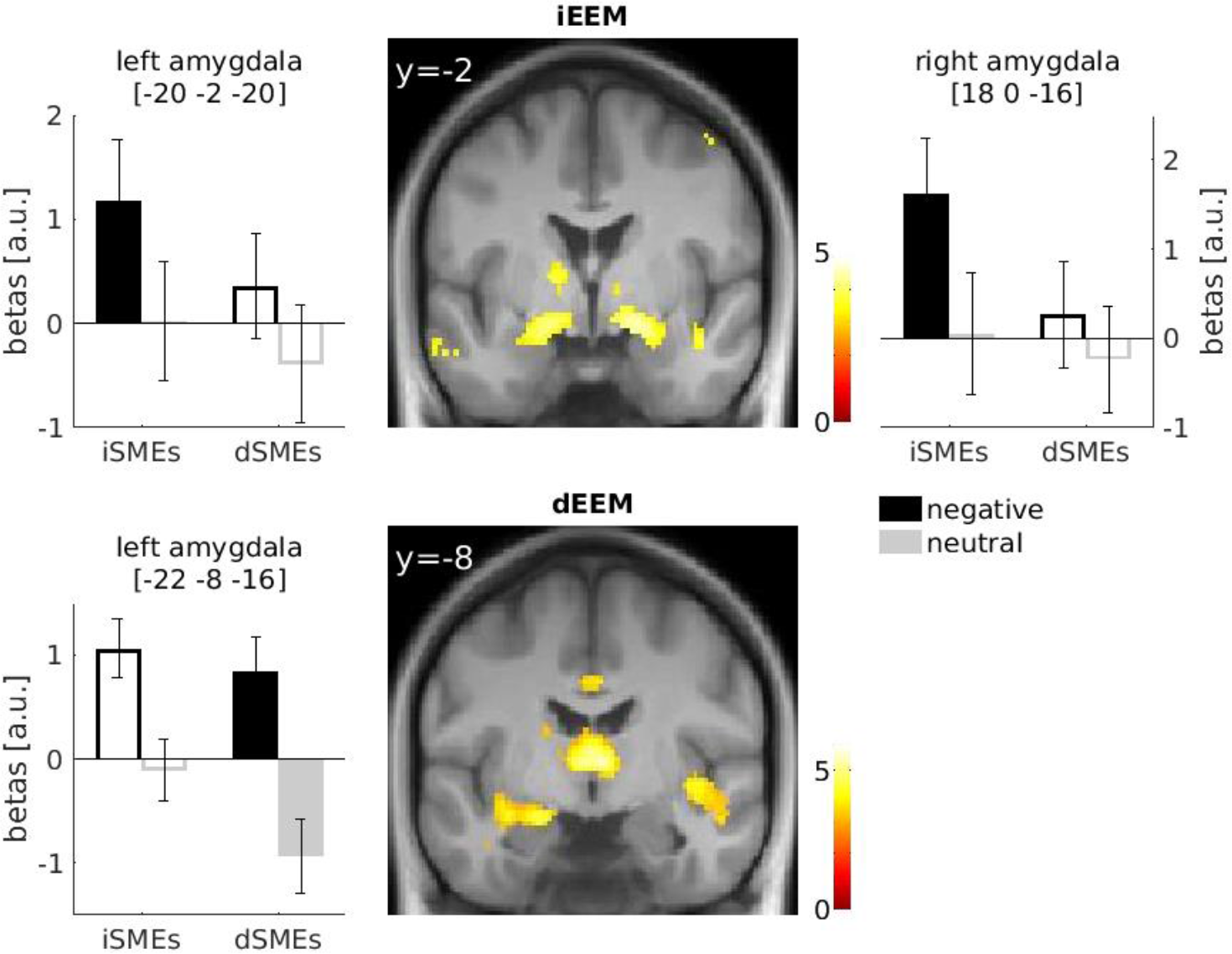
Amygdala activity associated with iEEM and dEEM. *Upper panel:* Activity in the bilateral amygdala correlated with iEEM. *Lower panel*: Activity in the left amygdala/anterior hippocampus correlated with dEEM. Negative and neutral subsequent memory effects (SMEs), i.e. activity during subsequent hits minus misses, for the immediate and delayed test are shown. The solid bars represent the effect (iEEM or dEEM) that was significant in the particular area, the contour bars the effects used as exclusive mask. Parameter estimates are shown in the peaks that survived the exclusive masking with the opposite contrast.

**Table 1:** Behavioral results in the immediate and delayed recognition tests. Familiarity and recollection were computed based on the dual process model of recognition memory (Koen et al. 2016, c response criterion, AUC area under the curve, mean and standard deviation).

|  | *hit rate* | *false alarm rate* | *c* | *d'* | *familiarity* | *recollection* | *AUC* |
| --- | --- | --- | --- | --- | --- | --- | --- |
| immediate test |  |  |  |  |  |  |  |
| negative | 0.85 (0.12) | 0.11 (0.09) | 0.17 (0.43) | 2.78 (1.26) | 1.63 (0.86) | 0.53 (0.29) | 0.92 (0.07) |
| neutral | 0.81 (0.13) | 0.13 (0.09) | 0.10 (0.50) | 2.42 (1.06) | 1.43 (0.66) | 0.46 (0.26) | 0.89 (0.08) |
| delayed test |  |  |  |  |  |  |  |
| negative | 0.68 (0.15) | 0.12 (0.09) | 0.43 (0.45) | 1.91 (0.80) | 1.32 (0.53) | 0.28 (0.21) | 0.84 (0.08) |
| neutral | 0.61 (0.19) | 0.17 (0.11) | 0.37 (0.43) | 1.38 (0.66) | 0.93 (0.51) | 0.20 (0.20) | 0.76 (0.11) |

The mean magnitude of both effects for participants was similar across the current study and when they participated in the previous large-scale behavioral study (Figure 1, left panel; GLM: main effect of study: F(1,68) = 0.07, p = 0.79; no interactions between the factors study, valence and delay, all p’s > 0.10). Most importantly, all post-hoc tests contrasting the same condition across studies, e.g. immediate corrected hit rate for neutral pictures, were not significant (all p’s > 0.40). Remarkably, the individual iEEM did not correlate across both studies (r = −0.01, p = 0.95) and the correlation was weak for dEEM (r = 0.23, p = 0.06). The correlation of the performance across studies within each of the four conditions was higher (immediate test: neutral r = 0.59, p < 0.0001, negative r = 0.40, p < 0.001; delayed test: neutral r = 0.39, p < 0.005, negative r = 0.42, p < 0.0005; see table 3 for the correlations of other memory measures).

Genotype had no effect on behavioral measures of iEEM of dEEM. A mixed GLM with delay and valence as within subject factors and genotype as between subject factor showed no effects for genotype and no genotype interaction for the adrenergic polymorphism (all ps > 0.31).

### fMRI Results

Various of the previously reported brain regions were associated with iEEM and dEEM, where overall activity in more areas was with dEEM (Table 2). Notably, activity in the bilateral amygdala was associated with iEEM whereas with dEEM only the left amygdala/anterior hippocampus correlated. This was confirmed by the exclusive masking procedure which revealed bilateral anterior amygdala for iEEM (xyz = [18 0 −16], Z = 4.39; xyz = [-20 −2 −20], Z = 3.56) and left posterior amygdala/anterior hippocampus for dEEM (xyz = [-26 −12 −14], Z = 4.04).

**Table 2:**
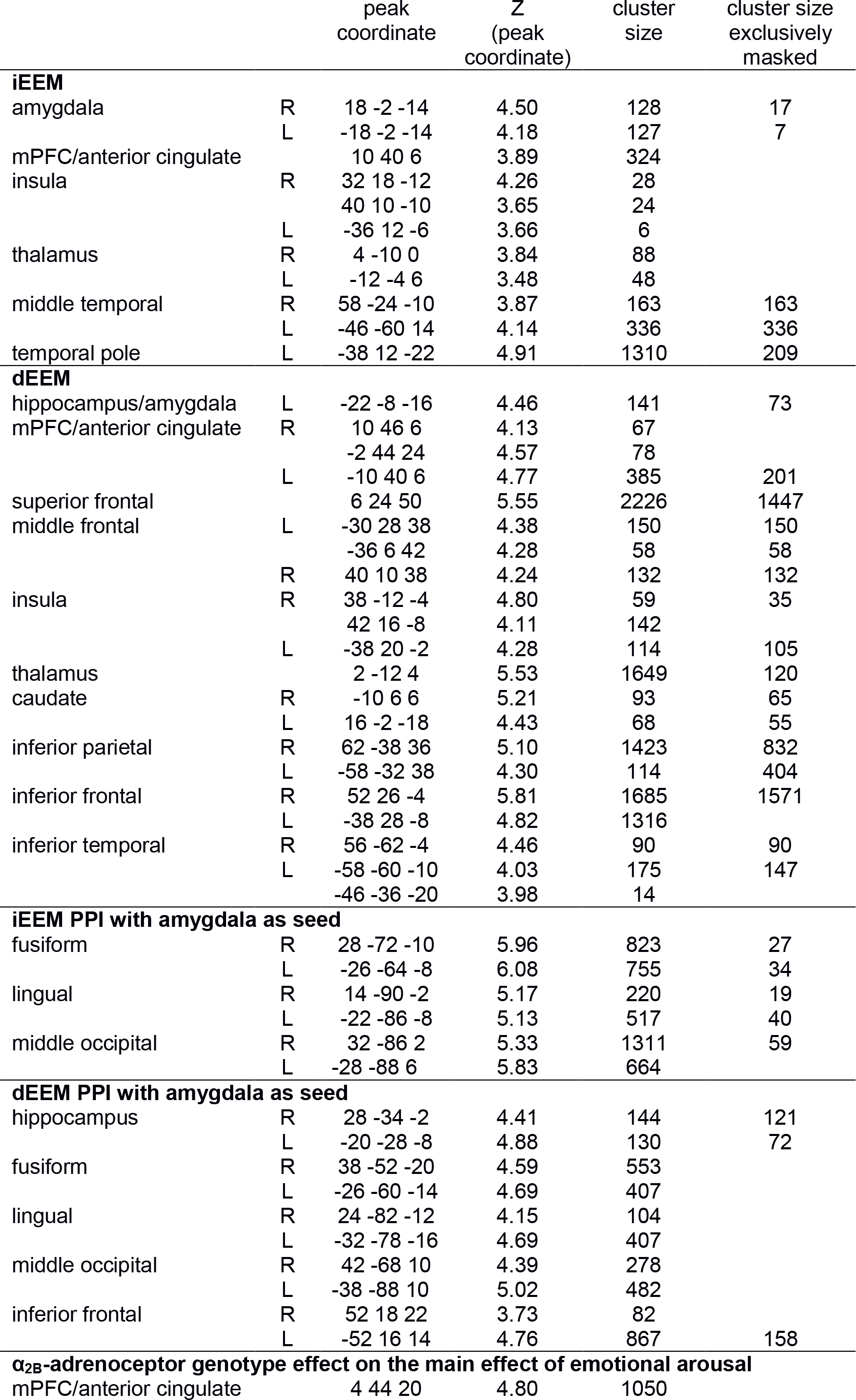
fMRI results. The number of voxels of activity clusters that survived the exclusive masking procedure with the opposite contrast are listed to show which clusters were predominantly related to iEEM or dEEM. (Cluster inducing threshold of p < 0.001 uncorrected)

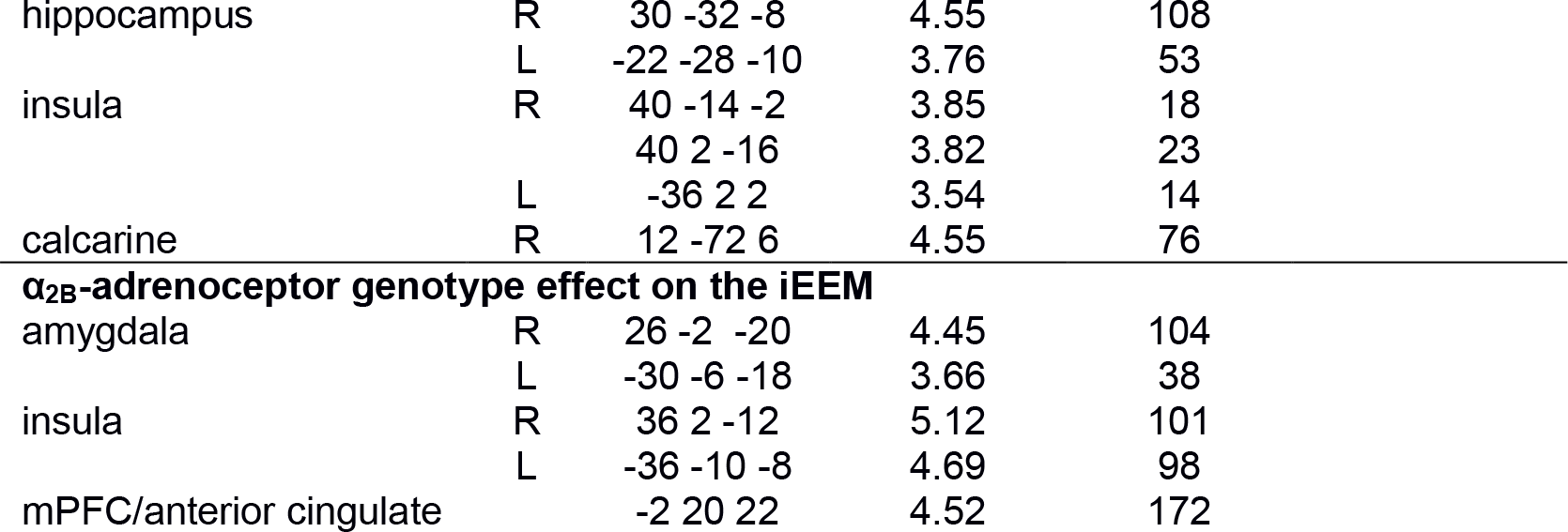

The PPI analyses using as seed a region in the left amygdala, identified by the main effect of emotion, revealed stronger coupling during successful encoding of negative than neutral pictures with large clusters in visual areas for iEEM (Figure 3, Table 2). During processing that was associated with more efficient consolidation for negative than neutral pictures the amygdala was coupled stronger with these visual areas and in addition the bilateral hippocampus and inferior frontal gyrus. The exclusive masking procedure revealed that clusters in visual areas, in particular in the fusiform (xyz = [28 −58 −6], Z = 4.59, xyz = [-26 −64 −4], Z = 5.17), lingual (xyz = [26 −64 −6], Z = 4.10, xyz = [-22 −88 0], Z = 4.85), and middle occipital gyrus (xyz = [22 −88 0], Z = 4.78) were predominantly related to iEEM whereas the hippocampus and inferior frontal gyrus to dEEM.

**Figure 3:**
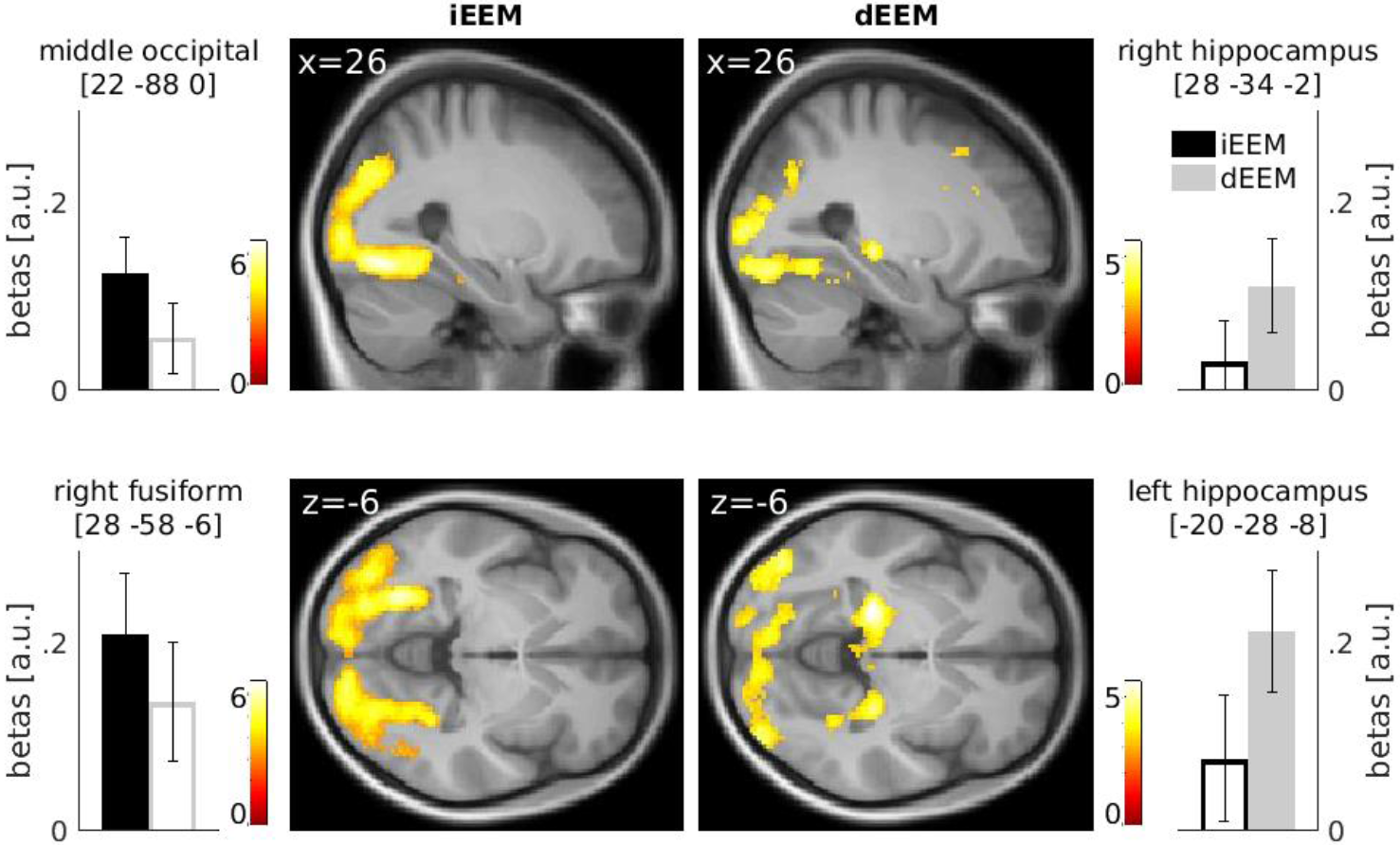
Amygdala coupling related to iEEM and dEEM. *Upper panel:* The amygdala was stronger coupled with the bilateral fusiform, lingual, and middle occipital gyri during successful encoding related to the iEEM. *Lower panel*: The amygdala was stronger coupled with the bilateral hippocampi, fusiform, lingual, middle occipital and inferior frontal during processing of negative than neutral pictures that were successfully consolidated. Parameter estimates are shown in the peaks that survived the exclusive masking with the opposite contrast. The solid bars represent the effect (iEEM or dEEM) that was significant in the particular area, the contour bars the effects used as exclusive mask.

The α_2B_-noradrenergic receptor genotype (32 wildtype, 37 deletion carriers) was associated with the greater activity in the medial PFC, bilateral hippocampus and visual cortex (Table 2) for the main effect of emotion, where deletion carriers showed greater activity in these areas for negative pictures. However, the largest p-value in the amygdala was only p = 0.29 (Z = 2.54) at xyz = [-30 −6 −28]. There were no brain areas associated with greater activity for participants with the wildtype of this adrenoceptor compared to the deletion carriers. However, wildtype carriers had greater iEEM-related activity in the bilateral amygdala (Figure 4, Table 2), where inclusive masking with brain areas related with iEEM, independent of genotype, resulted in greater associated activity in bilateral amygdala ([22 −4 −16], Z = 4.10; [-24 −4 −20], Z = 3.40), right insula ([40 10 −10], Z = 3.65), and right inferior frontal gyrus ([50 20 −8], Z = 4.19). Deletion carriers did not show any greater activity associated with iEEM and for dEEM neither group, i.e. deletion carriers or wildtype, showed significantly larger activity.

**Figure 4:**
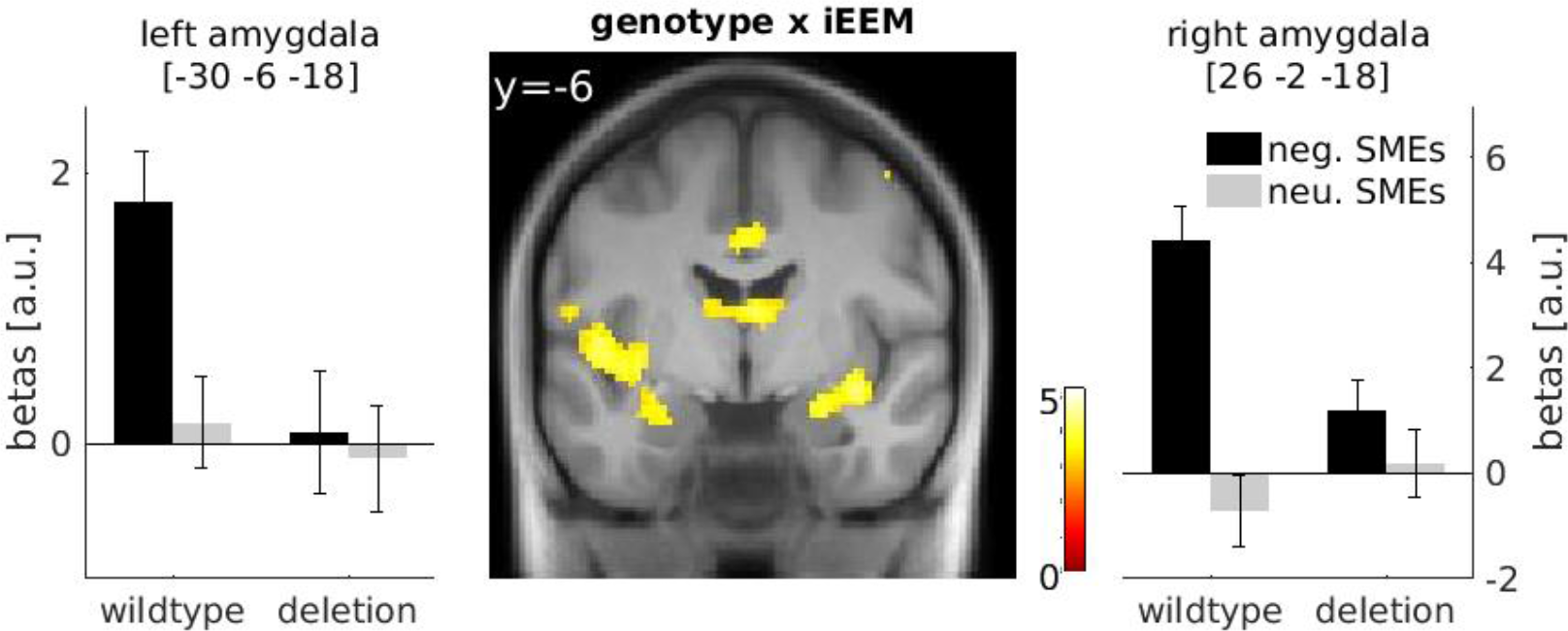
Interaction of α_2B_-adrenocepor genotype and amygdala activity associated with iEEM. Activity in the bilateral amygdala correlated stronger with iEEM in deletion carriers than α_2B_-adrenocepor wildtypes. Negative and neutral subsequent memory effects (SMEs) for the iEEM are shown for both genotypes groups

**Table 3:** Correlations of memory performance measures of the same 69 participants between the large-scale behavioral study 3.5 years ago and the current fMRI study. Rec - recollection, fam −familiarity according to the dual process model of recognition memory (Koen et al. 2016); EEM - Emotional Enhancement of Memory computed as differences between negative and neutral performance. ** p < 0.01; ^t^ p < 0.10.

| correlation behavioral study x fMRI study |  |  |  |  |  |  |  |  |
| --- | --- | --- | --- | --- | --- | --- | --- | --- |
|  | corrected<br>hit rate | hits | false<br>alarms | d' | c | fam | Rec | AUC |
| **immediate test** |  |  |  |  |  |  |  |  |
| negative | 0.40** | 0.37** | 0.33** | 0.33** | 0.20 | 0.03 | 0.19 | 0.37** |
| neutral | 0.59** | 0.55** | 0.58** | 0.41** | 0.41** | 0.14 | 0.35** | 0.58** |
| EEM | -0.01 | 0.13 | 0.15 | -0.13 | 0.16 | 0.07 | 0.18 | 0.10 |
| **delayed test** |  |  |  |  |  |  |  |  |
| negative | 0.42** | 0.36** | 0.50** | 0.42** | 0.34.** | 0.18 | -0.05 | 0.41** |
| neutral | 0.39** | 0.35** | 0.41** | 0.43** | 0.39** | 0.30* | 0.61** | 0.48** |
| EEM | 0.23<sup>†</sup> | 0.35** | 0.06 | 0.08 | 0.21 | -0.08 | 0.18 | 0.22 |

## Discussion

Recognition memory for negatively emotional pictures was better than for neutral pictures both immediately and after 24 hours, where this advantage increased after the consolidation delay, replicating previous findings, including from our previous large-scale behavioral study (Sharot et al. 2007; Sharot and Yonelinas 2008; Payne et al. 2008; Schümann et al. under review). Also similar to the previous findings from our large-scale behavioral and a smaller replication study, we found only a weak correlation between individual iEEM and dEEM (Schümann et al., under review). The consistently low correlation between iEEM and dEEM supports the hypothesis that both effects of emotional arousal, i.e. affect-biased attention and emotional synaptic tagging, rely at least partially on distinct neural substrates.

The mean iEEM and dEEM across participants of the current study was similar to the large-scale behavioral study about 3.5 years earlier, but only a weak correlation for individual dEEM across studies was significant. The two emotional memory paradigms differed in that in the current study, no emotionally positive pictures were included. Moreover, participants encoded in the MR scanner but retrieved on a computer outside of the scanner, which might have led to greater stress during encoding and to less context-dependent memory during recognition. The low correlations in individual iEEMs and dEEMs across the two studies might, therefore, be affected by such rather subtle differences in task and settings. However, these differences unlikely account for all of the individual variances; the low correlations might also suggest low retest reliability and/or stability of iEEM and dEEM. In other words, the typical paradigm used to assess the effects of emotional arousal on encoding and consolidation is either not well suited to measure potentially underlying personality traits, or iEEM and dEEM are not traits, but rather, states. Both interpretations are of relevance for behavior-genetic studies on EEM (Todd et al. 2011) and might explain why we previously failed to replicate a genetic influence of α_2B_-adrenoceptor polymorphism on iEEM (Schümann et al. under review). Importantly, previous studies reporting a influence of this polymorphism on iEEM used free recall (Rasch et al. 2009; de Quervain et al. 2007) where iEEM does not primarily depend on affect-biased attention during encoding but, to a larger degree, on processes that act during recall such as semantic relatedness and distinctiveness (Barnacle et al. 2016; Neely and Tse 2007; Talmi et al. 2017). The low retest reliability/stability of the EEMs is also more generally of interest because many behavior and/or imaging genetic studies use cognitive psychological measures, e.g. difference in performance between two conditions, with unknown psychometric characteristics as phenotype.

Our neuroimaging data suggest a lateralization of amygdala involvement in iEEM and dEEM. In particular, the right amygdala was correlated only with iEEM, the left amygdala with both iEEM and dEEM, and the left posterior amygdala/anterior hippocampus correlating only with dEEM. It is important to keep in mind that it cannot be ruled out that the overlap of both effects in the left amygdala might be caused by the unavoidable confound that many of the items contributing to dEEM also involved more affect-biased attention (iEEM) during encoding. The observed functional lateralization of the amygdala is consistent with previous reports of sex-dependent lateralization of dEEM. Specifically, the left amygdala contributes to consolidation only in women, the right in men (Cahill et al. 2001, 2004; Canli et al. 2002; Kilpatrick and Cahill 2003; Mackiewicz et al. 2006). Bilateral amygdala was associated with iEEM in both sexes (Mackiewicz et al. 2006). We did not find (with the given spatial resolution) evidence for a functional differentiation in the central, more dorsal, amygdala previously suggested to be involved in iEEM, and basolateral, more ventral, amygdala involved in dEEM (Mackiewicz et al. 2006). Instead, our data show an overlap of left amygdala activity associated with both effects, similar to other reports (Ritchey et al. 2008) and may suggest, if anything, an anterior-posterior dissociation.

The observed lateralized involvement of the amygdala in iEEM and dEEM would also be consistent with another proposed lateralization of amygdala function in terms of speed and duration of response to emotional stimuli. Specifically, a meta-analysis including a variety of stimuli (Sergerie et al. 2008), two face processing studies (Wright et al. 2001; Phillips et al. 2001) and also a study using emotional and neutral scenes (Kohno et al. 2015) have claimed that the right amygdala is part of a rapid emotional stimulus detection system whereas the left amygdala might be specialized for more sustained responses. The rapid detection of emotional stimuli could primarily reflect affect-biased attention, and the sustained response could be consistent with the modulation of emotional synaptic tagging. It is important to note that given that both sex-and function-dependent lateralizations of amygdala activity have been proposed, the lateralization of rapid vs. sustained effects might be different in men than in women. Besides sex and function, other processes leading to lateralization of amygdala activity have also been suggested (Glascher and Adolphs 2003). In line with a functional-and/or sex-dependent lateralization of amygdala activity, rodents show an anatomical lateralization of the amygdala which is in part sex-dependent (Johnson et al. 2008, 2012; Pfau et al. 2016).

Not only did activity in the amygdala differ between iEEM and dEEM, but its functional coupling with other cortical areas were also modulated depending on the EEM. In particular, the amygdala was functionally more strongly coupled with visual areas and the fusiform gyrus for iEEM and dEEM, and additionally with the hippocampus for dEEM. Both activity patterns overlapped, which might again be related to the confound that many pictures involved both affect-biased attention and emotional synaptic tagging. However, exclusive masking identified stronger coupling with fusiform and visual areas for iEEM and only with the hippocampus for dEEM. The stronger coupling with visual areas during successful encoding of negative than neutral pictures is consistent with known affect-biased attention effects on visual processing, in particular, the amygdala’s modulation of visual activity (Chen et al. 2014; Wendt et al. 2011). In contrast, dEEM was associated with greater coupling with the hippocampus, consistent with amygdala modulation of emotional synaptic tagging (McIntyre et al. 2012). Animal data also show that an electric stimulation of the amygdala after processing neutral objects results specifically in more effective consolidation via enhanced coupling with the hippocampus (Manns and Bass 2016). Similarly, the strength of connectivity of the amygdala with the parahippocampal gyrus during the processing of emotional pictures was associated with greater persistence of emotional memories over time (Ritchey et al. 2008).

Complementing the proposal that the relative independence of the effects of emotional arousal on iEEM and dEEM is due to differences in amygdala activity and coupling, on the neurobiological level, such independence can also be partly explained by the involvement of different adrenoceptor families or subtypes for each EEM (Hein 2006). The role of the α1- and β-adrenoceptor families, specifically in the effect of emotional arousal on memory consolidation, has been well established by a large body of pharmacological animal and human studies (Lonergan et al. 2013; McIntyre et al. 2012). However, the α2-adrenoceptor family might be predominantly involved in processes underlying affect-biased attention. The role of the α2-adrenoceptor family has been mainly studied using the α_2B_-adrenoceptor deletion polymorphism, which is associated with a less functional receptor and is moreover in complete linkage disequilibrium with a polymorphism in the promotor of the same gene that results in less transcription (Crassous et al. 2010; Nguyen et al. 2011). Deletion carriers show in one study greater amygdala activity and in another study greater activity in medial prefrontal as well as visual regions during processing of arousing stimuli (Rasch et al. 2009; Todd et al. 2011), an effect that we also found though not for the amygdala. Importantly, deletion carriers show enhanced affect-biased attention (Todd et al. 2013), greater iEEM in free recall (de Quervain et al. 2007) and enhanced recollection specifically for emotionally arousing items, which is consistent with more selective attention during encoding (Todd et al. 2014). In contrast, several studies reported no effect of α_2B_-adrenoceptor genotype on emotional memory consolidation (Naudts et al. 2012; Todd et al. 2014, 2015).

In the current and preceding large-scale behavioral study, we failed to replicate the larger iEEM in deletion carriers (as discussed above), but deletion carriers showed a smaller iEEM-related neural activity in the bilateral amygdala and insula. This neural activity difference for iEEM, in particular in the bilateral amygdala, supports the hypothesis that the α_2B_-adrenoceptor is involved in affect-biased attention but not emotional synaptic tagging. Moreover, it is consistent with our proposal based on the results of the whole sample that iEEM involves bilateral amygdala activity (Mackiewicz et al. 2006). However, at first sight, the direction of the difference in neural activity seems surprising because usually a behavioral effect such as larger free recall iEEM in deletion carriers is associated with greater neural activity. However, iEEM is the interaction of emotional and neutral SMEs which themselves reflect the increase in activity during encoding that is necessary for successful memory formation. The deletion polymorphism results in fewer and less functional α_2B_-adrenoceptors, which are at least partly expressed as presynaptic autoreceptors. Therefore, the deletion variant results in an increased noradrenaline release, which explains the greater neural activity with emotional processing we and others have observed. However, according to the Local Hot Spot model, increase in noradrenaline release results in an enhanced signal-to-noise ratio only at active synapses, which in turn increases encoding efficiency and thus suggests that a smaller activity increase is necessary for successful encoding (Mather et al. 2015).

Our replication of an effect of the α_2B_-adrenoceptor polymorphism on activity in the amygdala and other brain regions is also of a more general interest for a characterization of the role of this receptor subtype which was difficult because a selective antagonist was missing. Expression of this subtype was reported so far mostly in the periphery (e.g. vascular tissues) and centrally only in the spinal cord and thalamus. Consistent with this expression pattern, the phenotypes of α_2B_-adrenoceptor knockout mouse were described with respect to physiological parameters such as blood pressure (Bhalla et al. 2013). Only recently, it was shown that the α_2B_-adrenoceptor is more widely expressed in the brain and a role in gating and filtering of incoming sensory information has been suggested based on the behavioral phenotype of the knockout mice (Luhrs et al. 2016). The association of the α_2B_-adrenoceptor polymorphism with effect-biased attention related neural activity is somewhat supportive for this proposed function.

In conclusion, we replicated our previous behavioral finding that iEEM and dEEM where nearly uncorrelated across participants, suggesting these EEMs have distinct neural substrates. Although mean EEMs of the whole sample were of the same magnitude as 3.5 years ago, only dEEM was weakly correlated across time points, suggesting EEMs are not reliable or stable, at least when assessed by recognition tests. On the neural level, we observed a lateralized amygdala involvement. Specifically, whereas iEEM was associated with bilateral amygdala activity, dEEM was only associated with left and partly more posterior amygdala activity. This is consistent with previous reports of sex-and/or function-dependent amygdala lateralization. During successful immediate emotional memory formation, the amygdala was functionally more coupled with visual areas, consistent with affect-biased attentional modulation of visual processing. During successful delayed memory formation, the amygdala was more functionally coupled with the hippocampus, consistent with emotional synaptic tagging. The relatively lower iEEM-related activity in the bilateral amygdala in deletion carriers suggests that this receptor family is involved in affect-biased attention but not emotional synaptic tagging and is consistent with the proposal that bilateral amygdala is involved in such attention. Taken together, the current data support the hypothesis that the processes and underlying neural substrates of iEEM and dEEM are relatively independent of each other and suggest that the lateralized amygdala activity and its differential coupling with other cortical areas might be explained by different adrenoceptor families involved in the different EEMs.

## Acknowledgements

We would like to thank Gina Joue for her constructive comments that helped to improve the manuscript.

